## Supplemental Info for "Partitioning to ordered membrane domains regulates the kinetics of secretory traffic"

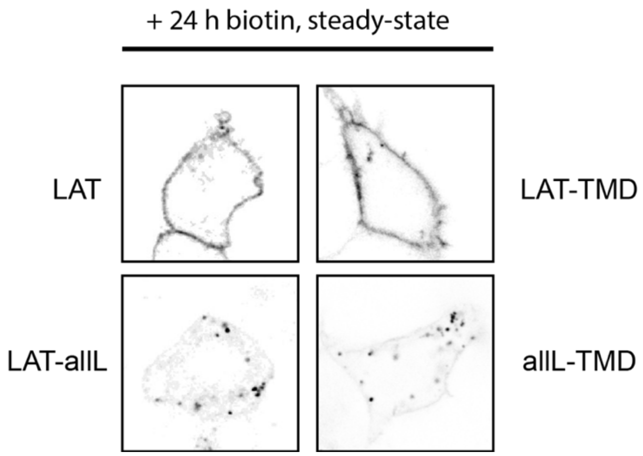


**Fig S1. Representative images of RUSH constructs after overnight (>10 hours) treatment with biotin.** Images show the steady-state distribution for these constructs: LAT and LAT-TMD accumulate at the PM, LAT-allL and allL-TMD accumulate in punctate structures previously identified as lysosomes.

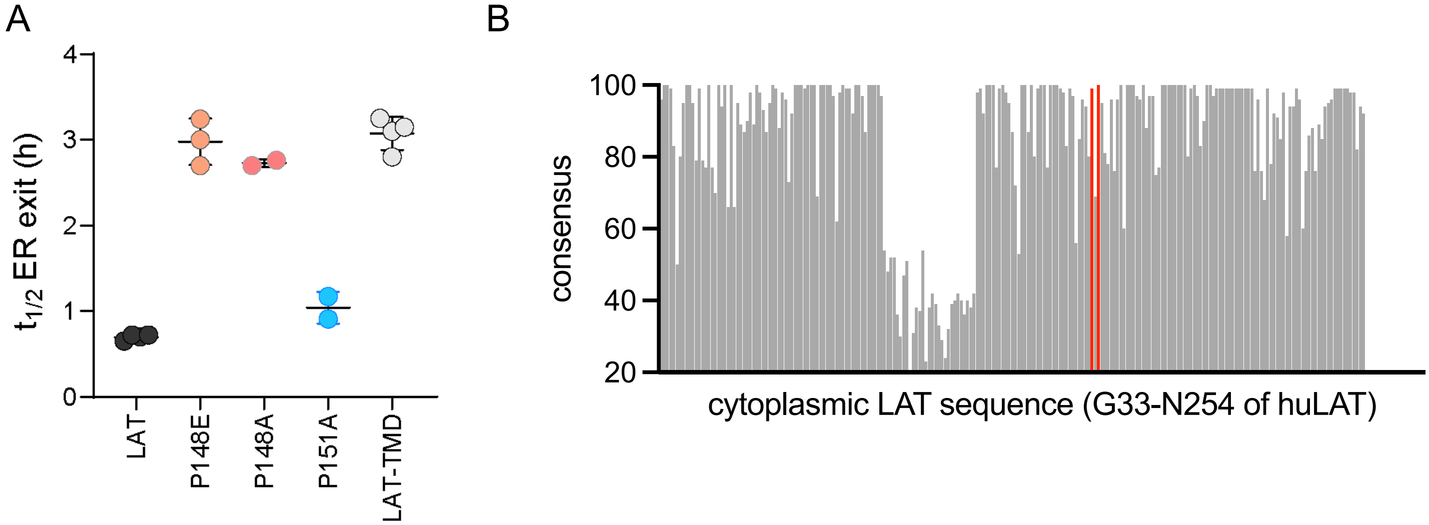


**Fig S2. COPII binding motif mediates fast ER exit of LAT.** (A) Half-time for ER exit for point mutations in the ΦxΦxΦ motif of LAT. P148 is critical for fast export from the ER, P151 is not. (B) Evolutionary conservation of cytoplasmic resides of LAT (denoted for residues G33 through the C-terminal N254 of human LAT). Red marks P148 and A150 for comparison.

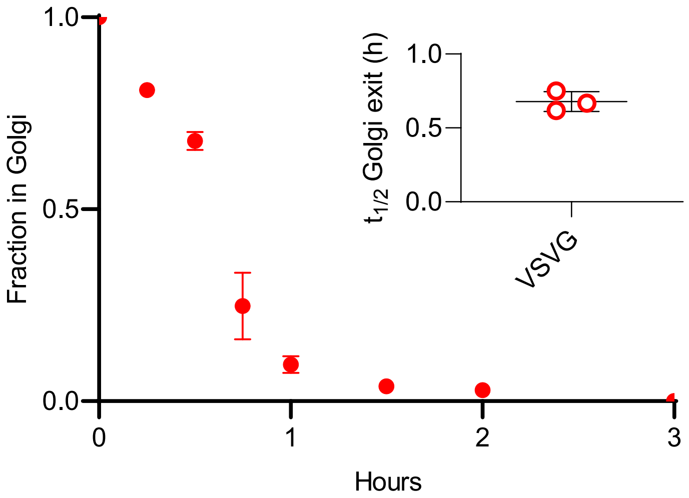


**Fig S3. Fraction of RUSH-VSVG in Golgi after biotin addition.** Inset represents repeats quantifications of the half-time of Golgi exit. Symbols represent average +/- st.dev. from 3 independent experiments.

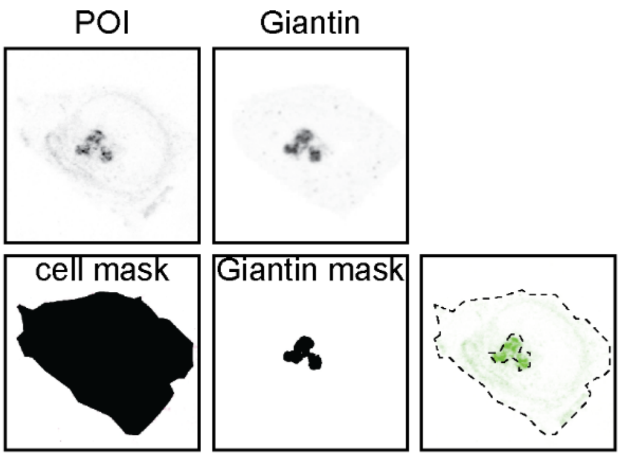


$$\text{Fraction in Golgi} = \frac{\left(\dfrac{\text{POI intensity inside giantin mask}}{\text{POI intensity inside cell mask}}\right)_t}{\left(\dfrac{\text{POI intensity inside giantin mask}}{\text{POI intensity inside cell mask}}\right)_{t=0}}$$

**Fig S4. Example of quantification of Golgi residence for protein of interest (POI).** Top panels are representative images, bottom panels are corresponding masks to calculate the fraction of POI in Golgi. Giantin was used as Golgi marker to create the mask for that organelle. Cells mask represents the cell border from the POI channel after background subtraction.

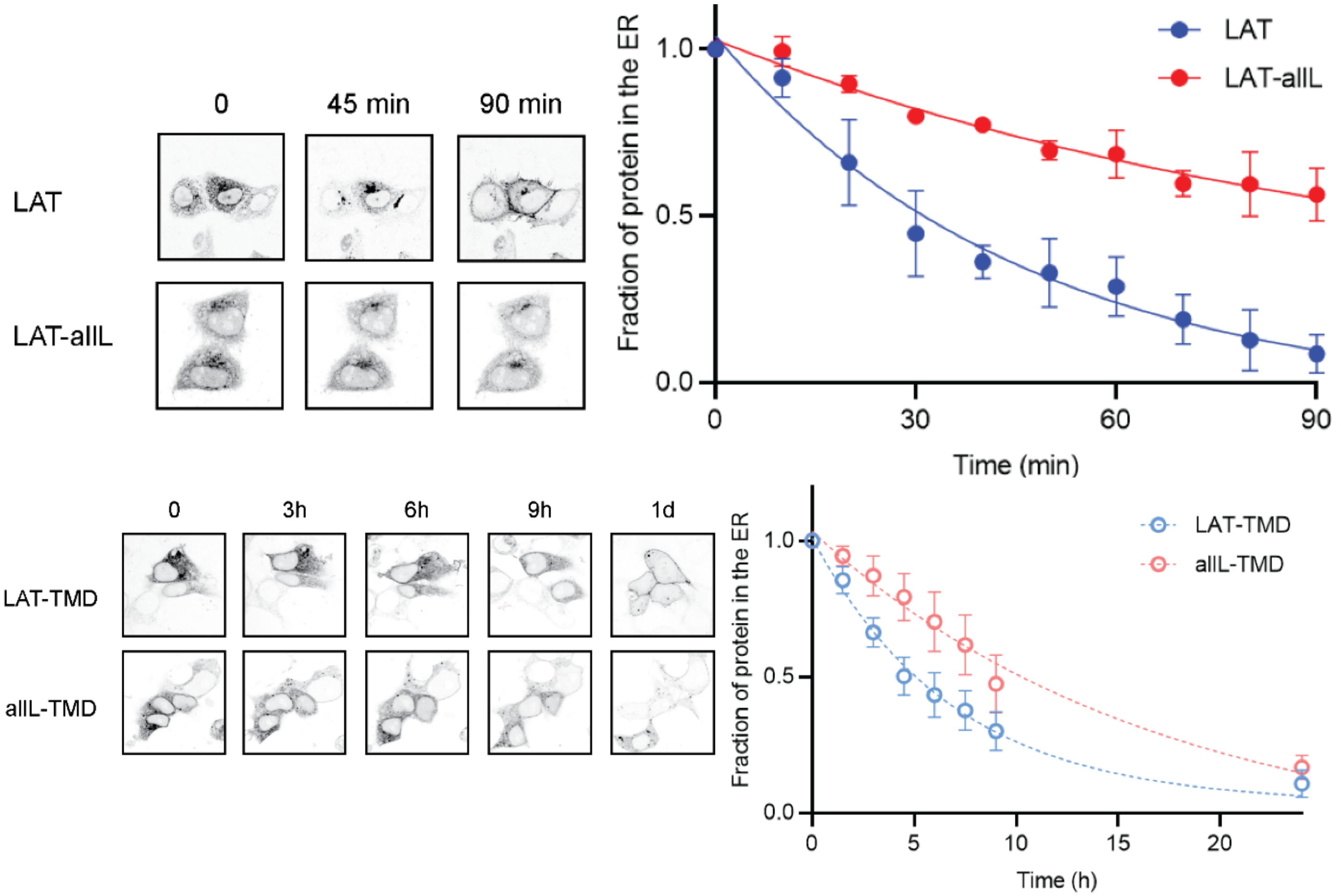


**Fig S5. Representative images of the experiments measuring ER exit kinetics within individual cells, used for the kinetic modeling.** Images show localization of RUSH constructs within a set of cells after biotin addition. Fraction in ER was quantified by making a mask of the protein at time 0 (i.e. before biotin addition) and calculating the remaining intensity within the mask (relative to total cellular intensity) at each subsequent time point. Top panels show full-length LAT and LAT-allL, bottom panels are LAT-TMD and allL-TMD. Symbols represent average +/- st.dev. from 3 independent experiments with multiple cells per experiment.

**Table S1. Raft affinity ($K_{p,raft}$) values for constructs used in this study.**

| **protein construct** | **$K_{p,raft}$ (mean±SD)** |
|---|---|
| GPI | 1.64 ± 0.09 |
| LAT | 1.33 ± 0.04 |
| LAT-TMD | 1.35 ± 0.06 |
| allA8L-TMD | 1.14 ± 0.13 |
| LAT-allL | 0.60 ± 0.07 |
| allL-TMD | 0.53 ± 0.01 |
| TfR | 0.37 ± 0.06 |
| LAX | 0.60 ± 0.02 |
| LAX-TMD | 0.76 ± 0.26 |
| VSVG | 0.50 ± 0.12 |
